# A highly sensitive protein-RNA cross-linking mass spectrometry workflow with enhanced structural modelling potential

**DOI:** 10.1101/2022.03.31.486537

**Authors:** Chris P. Sarnowski, Anna Knörlein, Tebbe de Vries, Michael Götze, Irene Beusch, Ruedi Aebersold, Frédéric H.-T. Allain, Jonathan Hall, Alexander Leitner

**Author notes:** These authors contributed equally (Anna Knörlein, Tebbe de Vries).

## Abstract

Protein-RNA interactions underpin many critical biological processes, demanding the development of technologies to precisely characterize their nature and functions. Many such technologies depend upon cross-linking under mild irradiation conditions to stabilize contacts between amino acids and nucleobases; for example, the cross-linking of stable isotope labelled RNA coupled to mass spectrometry (CLIR-MS) method. A deeper understanding of the CLIR-MS workflow is required to maximize its impact for structural biology, particularly addressing the low abundance of cross-linking products and the information content of spatial/geometric restraints reflected by a cross-link. Here, we present a vastly improved CLIR-MS pipeline that features enhanced sample preparation, data acquisition and interpretation. These advances significantly increase the number of detected cross-link products per sample. We demonstrate that the procedure is robust against variation of key experimental parameters, including irradiation energy and temperature. Using this improved protocol on four protein-RNA complexes representing canonical and non-canonical RNA-binding domains, we propose for the first time the distances encoded by protein-RNA cross-links, enabling their use as structural restraints. We also compared the cross-linking of canonical RNA with 4-thiouracil-labeled counterparts, showing slight, but noticeable differences. The improved understanding of protein-RNA cross-links refines the structural interpretation of complexes and facilitates the adoption of the method in integrative/hybrid structural biology.

## Introduction

Proteins and ribonucleic acids (RNAs) form functional units fundamental for cell functions(1–5). Understanding the structures of protein-RNA assemblies is crucial for understanding their respective functions. In cells, RNAs bind to a distinct set of common protein structure motifs, including the RNA recognition motifs(6) (RRMs), the zinc fingers(7) (ZnFs), and the DEAD box helicase domains(8) in ways that are generally structurally well understood. However, structural studies of prominent protein-RNA assemblies such as the ribosome(9, 10) and spliceosome(11, 12), as well as complexes that emerge from proteome-wide RNA-binding protein studies(13, 14), reveal a plethora of RNA-binding proteins that lack a recognizable canonical RNA binding domain.

As with RNA, proteins form functional complexes with other proteins. Cross-linking coupled with mass spectrometry (XL-MS) is a widely used approach in structural biology to reveal how protein complexes assemble and function. Specifically, protein-protein XL-MS establishes the spatial proximity between non-adjacent amino acids, thus providing information on which residues of two or more interacting proteins contact one another, or even how a single protein is folded(15–18). Despite the prevalence and importance of protein-RNA interactions, analogous XL-MS technology to study the structural and spatial nature of these interactions remains less mature than protein-protein XL-MS approaches.

A number of features of typical protein-protein XL-MS workflows(19) contribute to their practicality and prevalence. Firstly, protein samples are most commonly cross-linked using chemical reagents with known geometries (e.g. spacer lengths) and amino acid preferences, yielding residue-level localization and distance information for a given XL. Additionally, cross-linked peptides may be enriched for in samples, using techniques such as size-exclusion chromatography (SEC)(20) or immobilized metal affinity chromatography (IMAC)(21). Enrichment compensates to some extent for poor cross-linking efficencies, thus increasing the analytical coverage of cross-linked species. One of many specialized software tools(22) can then be used to interpret resulting mass spectra. Within each peptide, the precise site of amino acid cross-linking can be identified thanks to predictable peptide fragmentation during tandem mass spectrometry (MS/MS) analysis, and known amino acid-specific chemistry. Such workflows therefore reveal pairs of amino acids that reside in proximity with one another within the 3D structure of the folded protein(s), within a distance largely determined by the length of the chemical linker. These point-to-point distances can be specified as restraints in structural models, either as the sole source of experimental data(23), or combined with data from complementary structural techniques(24, 25), and provide a relatively fast way of analyzing protein structures in the solution state.

To utilize protein-RNA XL-MS data in a similar fashion, precise sites of XL attachment should be identifiable on both protein and RNA, at (or close to) single amino acid- or nucleotide-resolution, and the distance represented by such XLs needs to be well characterized(26, 27). Several experimental techniques have been developed that exploit UV cross-linking of proteins to nucleic acids(28), providing information about the XL with varying degrees of structural resolutions, ranging from whole molecule to single-residue. Techniques based on cross-linking and immunoprecipitation (CLIP)(29–33) provide localization of the binding site of a given protein on an RNA at high resolution using RNA sequencing, but the XL site on the protein remains mostly localized to a specific domain, providing insufficient detail for use as a restraint during structural modelling. Alternatively, some previously published MS based methods employ UV cross-linking to localize RNA contact sites at single amino acid positions of a protein(34–36), but only offer limited resolution on the RNA side of the interaction. While CLIP reliably revals the amino acids that mediate RNA interactions, it is of limited value in positioning the RNA sequence in relation to the protein and cannot therefore provide precise distance restraints for structural modelling pipelines. Our cross-linking of isotope-labelled RNA (CLIR-MS) workflow(37) overcomes this deficit and builds on previous protein-RNA XL-MS techniques(34, 38). Specifically, in a single experiment, CLIR-MS localizes the RNA cross-linking site, through stable isotope labelling of selected RNA segments to achieve up-to single nucleotide resolution, whilst simultaneously assigning the XL to a single amino acid on the interacting protein. The positional specificity afforded by this approach means protein-RNA XLs from a CLIR-MS experiment with position-specific RNA labelling represent ideal distance restraints in an analogous fashion to above-mentioned protein-protein XL-MS workflows.

In early investigations using CLIR-MS and related protein-RNA XL-MS workflows, the spatial/geometric properties reflected in a XL in the context of a protein-RNA complex were not systematically characterized. Thus, interpretations of protein-RNA cross-linking distances were reliant on unvalidated assumptions that such XLs occur at ‘zero-distance’(32), or on inferences from putative mechanisms for cross-linking chemistry based on prior literature(39–41). Hence, we hypothesized that a systematic study of cross-linking distance could provide more accurate interpretation of protein-RNA XLs, especially since there is much room for improved sensitivity in UV-mediated protein-RNA XL-MS experiments, due to low efficency of the UV cross-linking reactions. Often, less than 10% of starting material is converted to cross-linked complex using conventional conditions,(42) and only a minuscule fraction of this material – peptides at the RNA binding interface – is eventually analyzed. Improvements to sample preparation, data acquisition, and incorporation of a full set of well-characterized reaction products could together increase overall sensitivity of the CLIR-MS technique.

In this work we address these main limitations of the original CLIR-MS workflow. Specifically, we report enhancements to sample preparation and MS data acquisition, which collectively increase the numbers of XLs identified. Additionally, we take advantage of stable isotope labelling of RNA in CLIR-MS to characterize and validate the RNA-derived adducts to peptides more thoroughly, further enhancing the number of XLs. We then apply the optimized sample preparation and data analysis pipeline to a varied set of canonical RNA binding proteins (PTBP1, FOX1, and MBNL1) in complex with their cognate RNAs, and the non-canonical RNA binding ubiquitin-like domain of U2 snRNP protein SF3A1 in complex with stem-loop 4 (SL4) of the U1 snRNA. We compare XL identifications with published structures, and systematically assess the distance over which UV-induced protein-RNA XLs form. We also characterize the distinct properties of 4-thio-uracil compared with canonical uracil. Using these data, we show that CLIR-MS derived XLs alone contain sufficient information to describe the “occupancy space” of an RNA relative to a protein in complex. The work highlights the value of protein-RNA XL-MS data as a stand-alone data type, in addition to its previously demonstrated value as part of an integrative modelling pipeline(37), in understanding the vast emerging class of non-canonical RNA-binding proteins.

## Materials and Methods

### Protein expression and purification

PTBP1 and FOX1 RRM were prepared as described previously(37, 43). MBNL1 (amino acids 1-269 of MBNL140) was obtained in pGEX-6P1 (GE Healthcare). Plasmids were transformed into *E. coli* BL21 (DE3) codon+ (RIL) (Agilent Technologies) for protein expression. Cells were grown in K-MOPS minimal medium until OD_600 nm_ ∼0.5, shifted from 37 °C to 20 °C, and induced at OD_600 nm_ 0.7-0.8. Expression was carried out for 22-24 h. After harvesting the cells by centrifugation (15 min, 6000 rpm, 4 °C, Sorvall SLC6000 fixed angle rotor), dry pellets were frozen at −20 °C. Cells were thawed and resuspended to ∼0.25 g/mL in phosphate buffered saline (PBS; 140 mM NaCl, 2.7 mM KCl, 10 mM Na_2_HPO_4_, 1.8 mM KH_2_PO_4_ pH 7.3) using 1x cOmplete™ EDTA-free (Roche) tablet per 2 L culture. The cell suspension was homogenised using a 100 μm H10Z cell and Microfluidizer (Microfluidics) operated at 15000 psi, over three cycles. The lysate was subsequently clarified by centrifugation (60 min, 17000 rpm, 4 °C, Sorvall SS-34 fixed angle rotor). The resulting supernatant was incubated for 4-5 h at 4 °C on glutathione sepharose 4B (GE Healthcare) (3 mL per 2 L expression). After this, all further purification steps were performed at RT. The supernatant-resin slurry was loaded onto gravity flow columns and washed with 2 bed volumes PBS, followed by 10 bed volumes PBS + 1 M NaCl. Protein was eluted stepwise using 50 mM Tris, 500 mM NaCl, 50 mM glutathione (reduced) pH 8 (adjusted with NaOH). Pooled eluate was dialysed overnight at 4 °C into 20 mM sodium phosphate (NaP), 25 mM NaCl, 10 μM ZnSO_4_, 5 mM beta-mercaptoethanol pH 7. The GST-tag was cleaved by addition of HRV3C (1mg per 100 mg protein). Cleaved GST was separated by anion exchange chromatography (HiTrap Q HP 5 mL (GE Healthcare)). The equivalent of 1 L expression was injected per run, concentrated to 1 mL. To obtain non-degraded MBNL1Δ101 samples, flow through was again concentrated and subjected to gel-filtration chromatography, and buffer exchanged to 20 mM NaP, 50 mM NaCl, 10 μM ZnSO_4_, 5 mM beta-mercaptoethanol pH 6. Protein was then concentrated, aliquoted, snap frozen in liquid nitrogen, and stored at −80 °C until use.

For SF3A1-UBL (amino acids 704-793), the protein sequence fused to an N-terminal GB1 solubility tag and a 6x TEV-cleavable His-tag was cloned into pET24b (Novagen). Plasmids were transformed into *E. coli* BL21 (DE3) codon+ (RIL) (Agilent Technologies) for protein expression. Cells were induced at OD_600 nm_ 0.6-0.8 with 1 mM isopropyl-β-d-thiogalactopyransoide (IPTG). Expression was carried out for 4 h at 37 °C. Cells were grown in LB-medium (DIFCOTM LB-Broth, Fisher Scientific) with chloramphenicol and kanamycin. After harvesting the cells by centrifugation (10 min, 5000 x *g*, 4 °C, Sorvall SLC6000 fixed angle rotor), pellets were resuspended in 20 mM Tris pH 8, 1 M NaCl (buffer A), 10 mM imidazole, with cOmplete™ EDTA-free protease inhibitor. Cell lysis was carried out with a microfluidizer (Microfluidics), and the lysate centrifuged for clarification (30 min, 5000 x *g*, 4 °C). Protein purification was carried out by Ni-affinity chromatography, either with Ni-NTA beads (QIAGEN), stepwise by gravity flow, or using an ÄKTA Prime purification system (Amersham Biosciences) equipped with 5 mL HisTrap column (GE Healthcare), with an imidazole gradient of buffer B (20 mM Tris pH 8, 0.25 M NaCl, 500 mM imidazole). The buffer of the fusion proteins was exchanged by dialysis to buffer C (20 mM Tris pH 8, 0.25 M NaCl, 2.5 mM β-mercaptoethanol). The fusion protein was then cleaved overnight at 4 °C, using 6x His tag TEV (purified in house). GB1-6His and the His-TEV protease were removed from the solution with Ni-NTA beads, and the solution incubated with RNaseOUT (Invitrogen) for 15 min. The protein was then purified by size exclusion chromatography, using a HiLoad 16/60 Superdex 75 pg (GE) in 10 mM sodium phosphate pH 6, 50 mM NaCl. Protein was then concentrated, aliquoted, snap frozen in liquid nitrogen, and stored at −80 °C until use.

### Preparation of RNA

Multiple RNA isotope labelling strategies are available for CLIR-MS(44), and are employed in experiments presented here. ^13^C^15^N labelling results from transcription of RNA in isotopically labelled cell culture medium, as demonstrated previously(37). Alternatively, chemically synthesised short RNAs are employed, where RNA is synthesised using ^13^C ribonucleotides (also used elsewhere(40)). ^13^C^15^N *in vitro* transcribed RNA sequences, EMCV IRES RNA (sequence: 5’-GGAUACUGGCCGAAGCCGCUUGGAAUAAGGCCGGUGUGCGUUUGUCUAUAUGUUAUUUUCCACCAUAUUGCCGUCUUUUGGCAAUGUG-3’) and U1 snRNP SL4 RNA (sequence: 5’-GGGGACUGCGUUCGCGCUUUCCCC-3‘) were prepared as described previously(37). For chemically synthesised RNAs, standard phosphoramidites were purchased from Thermo Fisher Scientific. ^13^C ribose-labelled phosphoramidites were purchased from Pitsch Nucleic Acids. 4-thiouridine phosphoramidites were synthesised as described previously(45, 46). All other chemicals were obtained from Sigma-Aldrich, Fluorochem, TCI, and Fisher Scientific.

### Synthesis of oligonucleotides for model complexes

All oligonucleotides used were synthesised on a 50 nmol scale with the MM12 synthesiser (Bio Automation Inc.) using 500 Å UnyLinker CPG (Controlled-pore glass, ChemGenes) with standard synthesis conditions. Coupling time for the phosphoramidites was 2 × 180 s. The RNA phosphoramidites were used as 0.08 M solutions in dry acetonitrile (ACN). The activator BTT (CarboSynth) was prepared as 0.24 M solution in dry ACN. 0.02 M I_2_ solution in Tetrahydrofuran (THF)/Pyridine/water (70:20:10, v/v/v) was used as oxidising reagent. Capping reagent A was THF/lutidine/acetic anhydride (8:1:1) and capping reagent B was 16 % N-methylimidazole in THF. Detritylation was performed using 3 % dichloroacetic acid in dichloromethane.

For deprotection and cleavage from the solid support, the CPG was treated with gaseous methylamine for 1.5 h at 70 °C. For RNAs containing 4-thiouridine, the oligonucleotide was first incubated with 1 M DBU (1,8-diazabicyclo[5.4.0]undec-7-ene) in dry ACN (1 mL) for 3 h at RT and the CPG resin was washed with 5 mL in ACN. Afterwards the oligonucleotide was deprotected and cleaved from the solid support by using ammonia containing 50 mM NaSH (1 mL) at RT for 24 h. Desilylation for all RNA was carried out by treatment with a mixture of N-methyl-2-pyrrolidone (60 μL), triethylamine (30 μL), and triethylamine trihydrofluoride (40 μL) at 70 °C for 2 h. The reaction was quenched by adding trimethylethoxysilane (200 μL, 5 min, RT). Purification was carried out on an Agilent 1200 series preparative RP-HPLC using an XBridge OST C18 column (10 × 50 mm, 2.5 μm; Waters) at 65 °C with a flow rate of 5 mL/min, gradient 10–50 % B in 5 min (A= 0.1 M aqueous triethylamine/acetic acid, pH 8.0; B= 100 % ACN).

Fractions containing the DMT-protected product were collected, dried under vacuum, and treated with 40 % aqueous acetic acid for 15 min at RT to remove the DMT group. Samples were dried under vacuum and dissolved in 200 μL of water and purified by RP-HPLC on an XBridge OST C18 column (10 × 50 mm, 2.5 μm; Waters) at 65 °C with a flow rate of 5 mL/min, gradient 2–20 % B in 6 min (A= 0.1 M aqueous triethylamine/acetic acid, pH 8.0; B= 100 % ACN).

Fractions containing the desired product were collected and dried under vacuum. Mass and purity were confirmed by LC–MS (Agilent 1200/6130 system) on an Acquity OST C18 column (2.1 × 50 mm; Waters). The column oven was set to 65 °C, flowrate: 0.3 mL/min, gradient 1–35 % B in 15 min (A= water containing 0.4 M hexafluoroisopropanol, 15 mM triethylamine; B= methanol). UV absorption of the final products was measured on a NanoDrop 2000 spectrophotometer (Thermo Fisher Scientific).

### Cross-linking of protein-RNA complexes

5 nmol of purified protein-RNA complex was prepared per enrichment replicate, at concentration between 0.1 and 1.0 mg/mL, depending on the sample. At complex formation, the following protein:RNA stoichiometries were used: 1:1, 1:2, 1:4, 1:1, 1:1, for the FOX1, MBNL1, PTBP1-UCUCU, PTBP1-IRES (47), and SF3A1 complexes (48), respectively. RNA in the sample consisted of equimolar mixtures of unlabelled RNA and stable isotope labelled RNA (either ^13^C only for chemically synthesised RNA or ^13^C^15^N for in vitro transcribed RNA). The sample was subjected to 254 nm irradiation in a UVP Ultraviolet Crosslinker (Ultraviolet Products), with the sample cooled on a metal plate, pre-cooled to −20 °C, throughout. 4-thiouracil containing samples were irradiated four times with 150 mJ/cm^2^ at 365 nm in a Vilber Lourmat Bio-link BLX Crosslinker (Collegien). For the temperature and irradiation gradient experiments, sample cooling and irradiation were adjusted accordingly. Each irradiation step was followed by a pause of 1 min to allow the sample to cool. Unless otherwise stated, cross-linking irradiation energy was optimised using SDS-PAGE analysis to maximise cross-linking yield of the protein-RNA heterodimer, whilst minimising UV-induced multimers and degradation products.

### Digestion and enrichment

Samples were precipitated using 0.1 volumes 3 M sodium acetate (pH 5.2) and 3 volumes of ethanol precooled to −20 °C and kept at −20 °C for at least 2 h. Pellets of precipitated complexes were collected by centrifugation (30 min, 13000 × *g*, 4 °C). Pellets were washed in 2 volumes of 80 % ethanol in water (v/v) at −20 °C. The centrifugation step was repeated, and pellets air dried for 10 min. 50 µL of 50 mM Tris-HCl (pH 7.9) with 4 M urea was used to resuspend the pellet, and the solution then diluted with 150 µL 50 mM Tris-HCl, pH 7.9. 5 μg and 5 U per mg of cross-linked sample, of RNases A (Roche Diagnostics) and T1 (Thermo Scientific) respectively, were added, and RNA digestion carried out for 2 h at 52 °C on a ThermoMixer (Eppendorf). After cooling on ice, 2 μL of 1 M MgCl_2,_ and 125 U of benzonase (Sigma Aldrich) per mg of cross-linked complex, was added to each sample. Further RNA digestion was then carried out for 1 h at 37 °C on a ThermoMixer. Sequencing grade trypsin (Promega) was added at a 24:1 protein:enzyme ratio (w/w). Samples were incubated overnight at 37 °C on a shaking incubator, then heated to 70 °C for 10 min to deactivate trypsin. After deactivation, samples were cleaned up by solid-phase extraction (SepPak 50 mg tC18 cartridges, Waters), and dried in a vacuum centrifuge.

Titanium dioxide metal oxide affinity chromatography (MOAC) was used to enrich protein-RNA crosslinks as described previously(37),(49). In brief, dried samples were resuspended in 100 μL MOAC loading buffer (water:ACN:trifluoroacetic acid (TFA), 50:50:0.1 (v/v/v) with 300 mg/mL lactic acid), and incubated on a ThermoMixer at 1200 rpm for 30 min with 5 mg of pre-equilibrated TiO_2_ beads (10 μm Titansphere PhosTiO, GL Sciences). The beads were settled by centrifugation (1 min, 10000 × *g*, RT), and the supernatant carefully removed and discarded. 100 µL fresh MOAC loading buffer was added, and the sample incubated for a further 15 min. Centrifugation was repeated, the supernatant removed, and 100 µL MOAC washing buffer (water:ACN:TFA, 50:50:0.1 (v/v/v)) was added. After a further 15 min incubation, centrifugation was repeated, and the supernatant discarded. Peptide-RNA adducts were then eluted from the beads with 50 µL MOAC elution buffer (50 mM ammonium phosphate, pH 10.5). Samples were incubated for 15 min, and beads again settled by centrifugation. The supernatant was carefully collected, stored on ice, and elution repeated a second time and combined with the first eluate. The eluate solution was immediately acidified to pH 2-3 with TFA. Eluates were purified with C_18_ solid phase extraction using self-packed Stage tips. In brief, two layers of C_18_ membrane (Empore, 3M) packed in a 200 µL tip (MaxRecovery, Axygen) were washed with 80 μL 100% ACN with 0.1% formic acid (FA), 80% ACN with 0.1% FA in water, then equilibrated twice with 80 μL 5% ACN with 0.1% FA in water. Each sample was applied to the membrane, and the membrane then washed 3 times with 80 μL 5% ACN with 0.1% FA in water. Purified peptide-RNA adducts were eluted from the membrane three times with 50 μL 50% ACN with 0.1% FA in water. LoBind tubes (Eppendorf) pre-washed with 50% ACN with 0.1% FA in water were used to collect purified peptide-RNA adducts. The sample was then dried in a vacuum centrifuge. For PTBP1-IRES samples cleaned up with C_18_ cartridges in **Figure 1**, SepPak tC18 cartridges (Waters) were used for clean-up instead.

**Figure 1:**
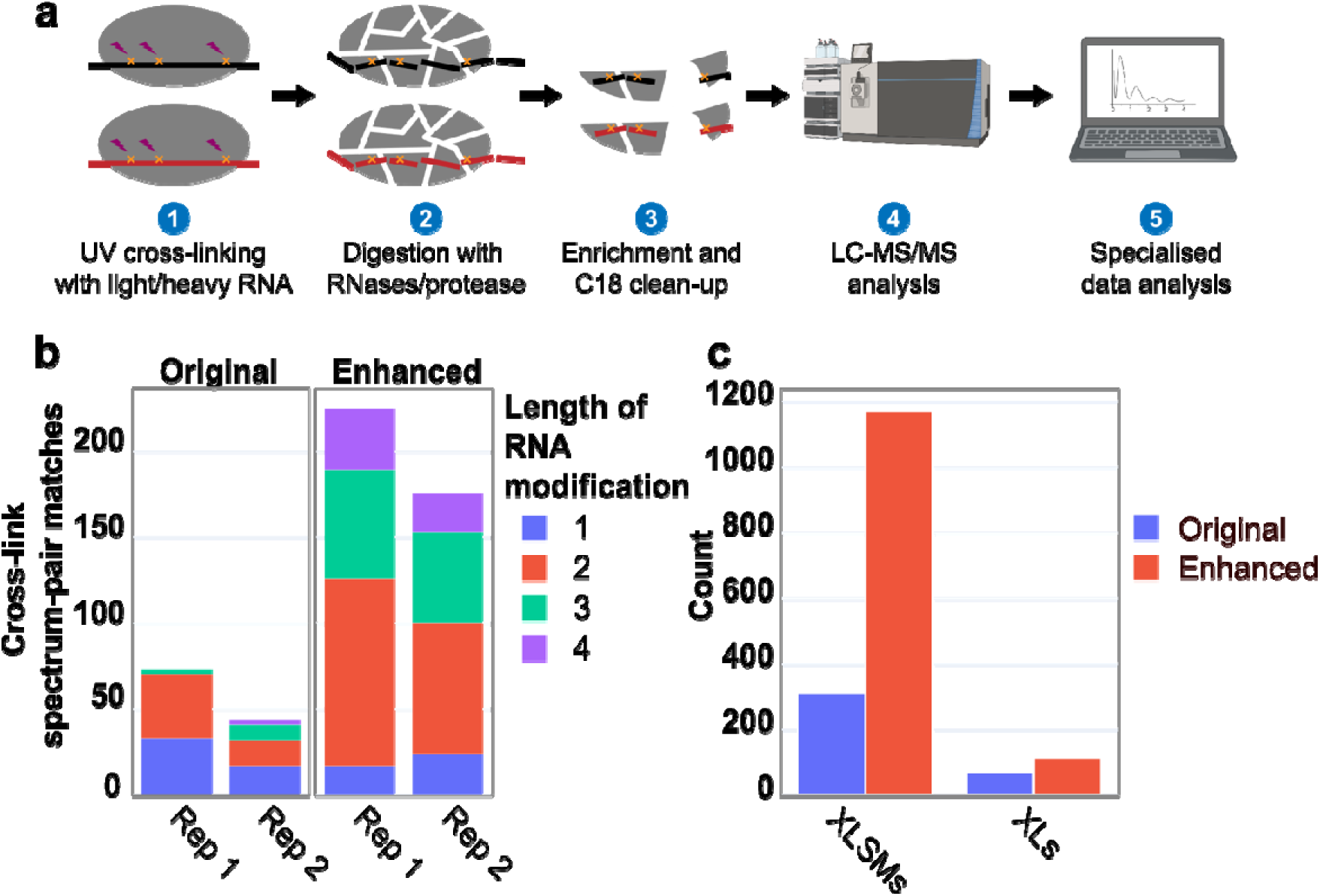
Optimization of sample preparation, data acquisition, and data analysis for the CLIR-MS workflow. a) Overview of the CLIR-MS workflow, as established in Dorn *et al* (37). b) Numbers of identifications produced when the FOX1-FBE complex is prepared and analyzed with and without the enhancements in method design. The optimizations from the PTBP1-IRES complex transfer to other protein-RNA complexes. Rep = Replicate. c) Comparison of xQuest searches of the same FOX1-FBE data (UV cross-linked condition only) using the expanded set of cross-linking products identified in this work, in comparison to the more restrictive set of modifications specified in previous CLIR-MS work (37).

### Analysis of samples with LC-MS/MS

Each dried sample was resuspended in 20 µL mobile phase A (described below), and 5 µL of each sample was injected for LC-MS/MS analysis. For MS method optimisation experiments in **Figure 1**, resulting samples from multiple enrichment replicates were pooled, and 3 μL samples were injected to evaluate each acquisition method. For data shown in all figures LC-MS/MS analysis was performed using an Easy-nLC 1200 HPLC system (Thermo Fisher Scientific) coupled to an Orbitrap Fusion Lumos mass spectrometer (Thermo Fisher Scientific) equipped with a Nanoflex (Thermo Fisher Scientific) nanoflow electrospray source. Peptide-RNA adducts were separated using a PepMap RSLC column (250 mm × 75 um, 2 µm particle size, Thermo Fisher Scientific) with gradient of 6-40% mobile phase B (A= water:ACN:FA, 98:2:0.15 (v/v/v); B= water:ACN:FA acid, 20:80:0.15 (v/v/v)) with a flow rate of 300 nL/min over 60 min.

The Orbitrap Fusion Lumos was used in data dependent acquisition mode, and the Orbitrap mass analyser used for precursor ion spectra acquisition, with resolution of 120000. Fragmentation was achieved with higher-energy collisional dissociation (HCD), using stepped collision energies of 21.85 %, 23 % and 24.15 %. Precursor ions with charge states between +2 and +7 were selected with a quadrupole isolation window of 1.2 m/z and cycle time of 3 s, with a dynamic exclusion period of 30 s. Resultant fragment ions were detected in the ion trap at rapid resolution. For experiments related to the SF3A1-SL4 complex, fragment ions were detected in the Orbitrap using a resolution of 30000.

### Data analysis with xQuest (light-heavy labelled species)

Data files produced by the mass spectrometer (Thermo Fisher .RAW format) were converted to centroided mzXML files using msconvert.exe (ProteoWizard msConvert v.3.0.9393c(50)). Files were then searched using xQuest (version 2.1.5, available at https://gitlab.ethz.ch/leitner_lab/xquest_xprophet)(51, 52) against a database containing only the sequence of the target protein. xQuest was originally designed to analyse protein-protein XL-MS data, with workflows in which an equimolar mixture of light and heavy isotopes of a chemical cross-linking reagent have been used to covalently link peptides. During a CLIR-MS experiment, a light-heavy stable isotope labelled RNA segment cross-linked to a peptide behaves similarly to a monolink(18) (type 0 cross-link(53)) in peptide-peptide cross-linking nomenclature, thus enabling xQuest to also process such data. Scripts from the Python package *RNxQuest* were used to define these searches, consolidate the resulting outputs(54).

All amino acid types were permitted as possible modification sites, and all possible RNA-derived adducts of 1-4 nucleotides in length, based on the RNA sequence of the respective complex and including all loss products, were considered possible modifications. For RNA produced by solid phase synthesis, a delta mass of 5.016774 Da per labelled nucleotide in the expected RNA modification was specified, to restrict identifications to those containing labelled RNA. For *in vitro* transcribed sequences, delta masses were defined according to expected ^13^C^15^N labelling patterns, described previously(37). A +/- 15 ppm mass tolerance window and 60 s retention time tolerance were used for pairing of light-heavy species. Further parameters for xQuest searching (described previously(52)): Enzyme = trypsin, maximum missed cleavages = 2, MS1 mass tolerance = 10 ppm, MS2 mass tolerance = 0.2 Da for ion trap MS2 data or 10 ppm for Orbitrap MS2 data. Identifications with an ld.Score > 20 (according to the scoring scheme described previously(52)) were considered. In this work, we considered robust identifications as particularly important for method development. Given that the ld.Score reflects spectral quality, use of a higher score threshold than that representing a false discovery rate of <1% ensures additional stringency in favor high quality spectral identifications. However, as described elsewhere(54), we do not consider this necessary in more routine applications of the technology. Further processing was completed using custom Python 3.7.1 scripts, and functions from the *RNxQuest* Python package(54). CLIR-MS plots shown here have amino acid numbering retrospectively adjusted from the FASTA file numbering to match prior structural models of each complex published in the PDB. Identifications were further refined for mass accuracy. Where multiple identifications were produced against the same spectrum, only the highest scoring identification was retained. Raw data files and xQuest search engine result files are accessible in PRIDE, described in ‘Data Availability’.

The sparse nature of metal oxide-enriched protein-RNA adduct samples means XLSMs made are often near the limits of detection by MS, and hence are vulnerable to fluctuations in instrument performance over time. Comparisons are therefore only made within batches where data acquisition took place at a similar time.

### Data analysis with MSFragger (open modification search)

Thermo Fisher .RAW files were converted as above. Default parameters for an open search using MSFragger (v2.1) were loaded, and the following modified: modification range = 150-1400 Da, fragment mass tolerance = 0.2 Da, allowed missed cleavages = 2, minimum peptide length = 5, top peaks = 250, min_fragments_modelling = 3, min_matched_fragments = 5, allow_multiple_variable_mods_on_residue = 1. The search was executed using the FragPipe GUI (v11.0), and results output to a comma-separated value (csv) file. Identifications with an expect score of greater than 0.05 were removed. Remaining matches to decoy sequences were also removed. Mass additions to peptides were then aggregated in 0.1 Da mass bins, and RNA-derived mass additions to peptides were manually annotated according to existing literature where possible, to form a putative list of RNA-derived products. Putative products were subjected to validation using a light-heavy dependent xQuest search.

### Comparison of CLIR-MS results with published structures

Protein-RNA XLs identified from the xQuest search were filtered, retaining only identifications where a mononucleotide RNA was cross-linked to a peptide. A custom script in PyMOL (version 2.3.2, Schrödinger) was used to compare identified XLs with published ensembles of models derived from NMR spectroscopy for each model complex; distances for XLs were measured from the Cα atom of the amino acid position in the XL identification, to N1 (pyrimidines) or N9 (purines) of the closest nucleotide in the structure matching the nucleotide type observed in the XL identification. Distances were outputted to a csv file and distributions plotted using the Python package Plotly. Identified protein-RNA XLs were plotted on published structures using PyXlinkViewer(55), with code modified to plot nucleotide XLs.

### Visualization of structural information content with DisVis

A restraint list of amino acid and nucleotide positions was prepared from protein-RNA XLs identified by CLIR-MS. Amino acid positions were taken directly from the xQuest results. For complexes in **Figure 4a-c**, RNA positions were selected based on the closest matching nucleotide, as measured in published structures. For the U1 snRNA SL4 sequence in **Figure 4d**, RNA positions were derived by overlaying non-redundant RNA compositions detected at every cross-linked amino acid position with the full RNA sequence. The 5% highest scoring positions were used to define RNA sequence positions for restraints. In all complexes shown in **Figure 4**, the restraint distances were defined between a minimum of 0 Å, and a maximum of 12 Å, with 12 Å corresponding to the upper quartile of all cross-linking distances measured against published structural ensembles of canonical RNA binding proteins in **Figure 3e-g**. Restraints were specified from Cα of the amino acid position to N1 (pyrimidines) or N9 (purines) of the nucleotide position. For complexes in **Figure 4a-c**, published structures were downloaded from the PDB, and protein and RNA chains were exported as separate molecules. For the complex in **Figure 4d**, standalone protein and RNA structures available in the PDB were downloaded separately (PDB ID: 1ZKH for the unbound SF3A1 UBL structure(56), and PDB ID: 6QX9 for the structured SL4 RNA(57)). Protein structures were specified as fixed chains, and RNA structures as scanning chains. Individual protein and RNA chains submitted to the DisVis web server(58, 59) (https://wenmr.science.uu.nl/disvis/), together with the restraints file in the required format. “Occupancy Analysis” was enabled, and “Complete Scanning” was selected. All other parameters were left at default values. Outputs from DisVis analysis were visualised with UCSF Chimera(60).

## Results

### An optimized CLIR-MS protocol with increased sensitivity

The CLIR-MS sample preparation protocol (**Figure 1a**) comprises cross-linking of protein-RNA complexes by UV irradiation, digestion of complexes with nucleases and proteases, enrichment for cross-linked peptide-RNA adducts with titanium oxide metal oxide affinity chromatography (MOAC), sample clean-up using C_18_ solid phase extraction (SPE), and analysis by liquid chromatography coupled to tandem mass spectrometry (LC-MS/MS)(37). To improve the sensitivity of the CLIR-MS method, we systematically optimized these steps to improve recovery and detection of protein-RNA adducts. We assumed that this would result in both increased total numbers of assigned MS/MS spectra (cross-link spectrum matches, XLSMs) during data analysis, as well as an increased number of unique peptide-RNA identifications from these XLSMs. Optimizations were carried out using the previously studied polypyrimidine tract binding protein 1 in complex with the internal ribosome entry site of the encephalomyocarditis virus (PTBP1-IRES complex). Multiple RNA binding domains, and the relatively long bound RNA (88 nucleotides) provide many interaction regions, resulting in a rich variety of cross-linked peptide-RNA adducts. In brief, both a miniaturization of the sample clean-up step and a switch from collision-induced dissociation (CID) in the ion trap to higher-energy CID (HCD) contributed to the performance increase. The improvements were not only reflected in the number of XLSMs (increased from 117 to 401), but also in the increased number of unique peptide sequences (increased from 8 to 9), cross-link sites (increased from 18 to 23), RNA adduct compositions increased from 8 to 11), and unique cross-link products (non-redundant combination of peptide sequence, RNA adduct, and cross-link site, increased from 42 to 114).

To test whether the optimizations made using the PTBP1-IRES complex could be generalized, we compared our original and revised protocols on a different protein-RNA complex, the FOX1 RRM with its cognate RNA, the FOX Binding Element (5’-UGCAUGU-3’, FBE). FOX1 regulates eukaryotic RNA splicing by binding the RNA sequence UGCAUG(62), and is structurally well characterized (PDB ID: 2ERR) using solution state NMR spectroscopy(43), meaning that identified XLs could be validated against a known structure. The number of XLSMs increased from approximately 50 per replicate with the previous protocol to around 200 per replicate with the enhanced protocol (mean of both replicates, **Figure 1b**), suggesting that optimizations made using the PTBP1-IRES complex have broad applicability.

Using the improvements described above and the FOX1-FBE sample, we next investigated how conditions during the UV cross-linking reaction (**Figure 1a** step 1) could impact the cross-linked species detected in a CLIR-MS experiment. We evaluated the impact of cross-linking temperature as well as UV irradiation energy dosage on the XLs obtained (data not shown). We found that even under both perturbations, obtained XLs remain qualitatively consistent, suggesting that unique structural information obtained in a CLIR-MS experiment is highly robust against procedural variation.

In summary, with respect to sample preparation and data acquisition, we found that miniaturization of sample clean-up and optimization of LC-MS/MS fragmentation conditions contributed to an increase in the number of XLSMs made in a CLIR-MS experiment, that these enhancements are generalizable beyond a single protein-RNA complex, and that the qualitative XL identifications made in a CLIR-MS experiment remained consistent over a broad range of UV cross-linking reaction conditions.

### Characterization of new cross-link products enhance CLIR-MS coverage

Next, we attempted a more thorough characterization of UV-induced RNA-derived peptide modifications found in a CLIR-MS sample, to investigate whether a broader set of reaction products could be routinely incorporated into the data analysis strategy (**Figure 1a** step 5). Stable-isotope labelled RNA is primarily employed to localize XLs to specific nucleotides in the RNA – those which are present in both light and heavy form in the sample result in a characteristic mass shift. Additionally, the presence of these mass-shifted species confirms that peptide modifications truly derive from RNA, rather than any potential side product of the UV cross-linking reaction with coincident mass. In the present work, two labelling schemes, either with *in vitro* transcribed RNA incorporating ^13^C and ^15^N, or chemical synthesis of RNA using nucleotides containing ^13^C ribose, were used (further details in **Materials and Methods**). Many different RNA-derived products have been described in previous peptide-centric mass spectrometric analyses of UV-induced protein-RNA XLs(34, 36, 38, 63, 64). These suggest that a variety of neutral losses, including those corresponding to atomic compositions such as -H_2_, -H_2_O, -HPO_3_, occur from the peptide-RNA adduct during cross-linking, sample preparation, or mass spectrometric analysis. To better explore these RNA moieties, a new FOX1-FBE dataset with different replicate structure was generated, comprising both a UV cross-linked condition, and a non-cross-linked control. Data were searched using an open-modification approach with the MSFragger search engine(65), and a putative enlarged list of RNA adduct types was discovered. Resulting mass shifts were assigned identities where possible, according to prior literature (**Table 1**). The list of putative RNA adduct types was then validated using the properties provided by light-heavy isotope paired RNA, and the xQuest search engine, using the RNxQuest companion package(54).

**Table 1:**
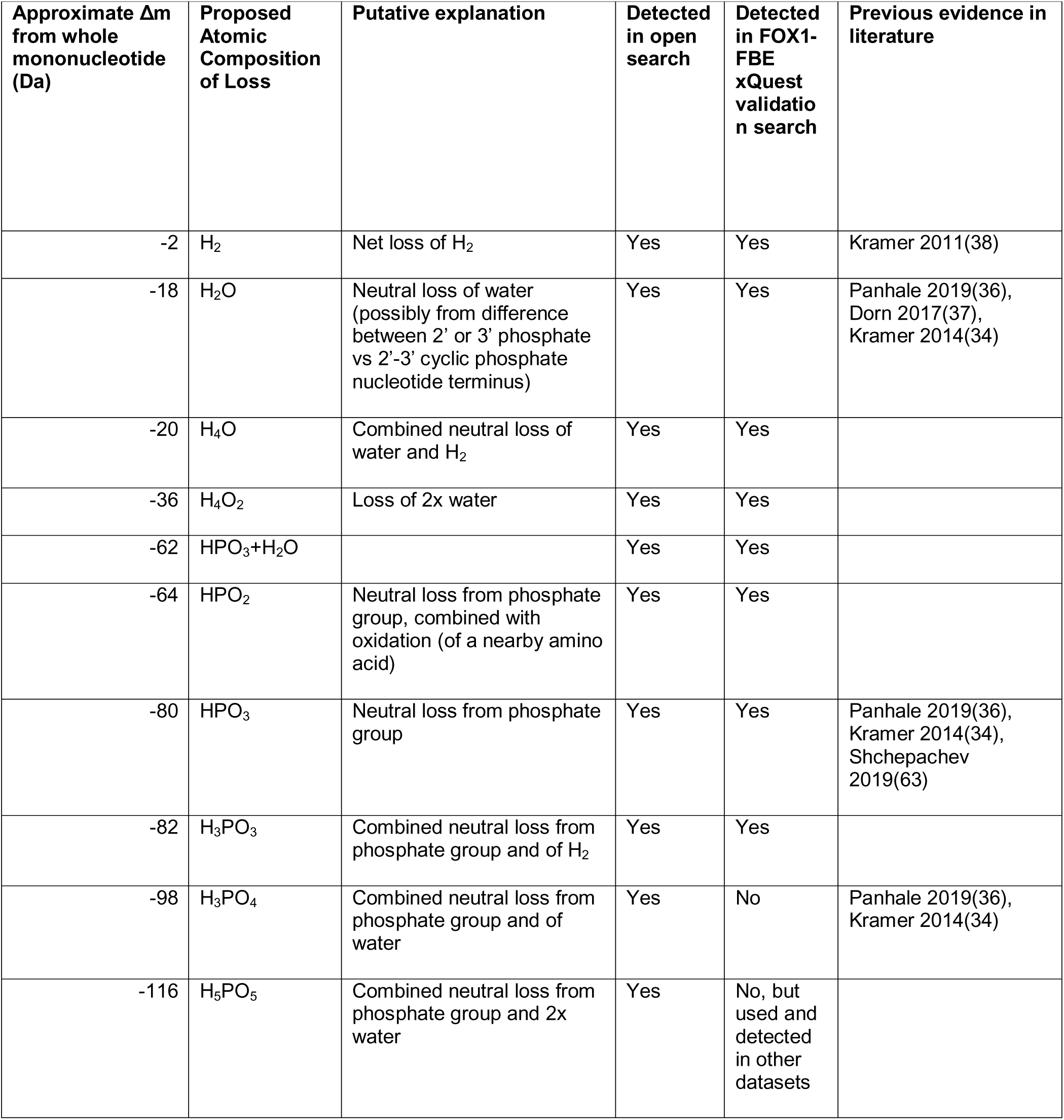
Types of RNA-derived peptide adducts validated for use in further xQuest searches. Losses from RNA-derived peptide adducts that were putatively detected in an open modification search of a CLIR-MS data set generated using the FOX1-FBE complex, and subsequently validated by xQuest search, specifying such losses in search parameters. Only adduct types that were validated by xQuest search in the FOX1-FBE complex were used for further xQuest searches.

Product types subjected to confirmation using light-heavy isotope xQuest search are shown in **Table 1**. Based on these results, we used an expanded list of RNA adducts for subsequent searches – these are listed as “validated” in **Table 1** and have been subsequently used in other recent work by our group(66). Whilst the precise mechanism of production of this full array of adducts requires further investigation, their existence in both light and heavy isotopic form suggests these represent true protein-RNA XL products.

To conclusively assess the impact of an expanded neutral loss product set on the total number of protein-RNA XL identifications made in a CLIR-MS experiment, the original and expanded lists of neutral loss products were used to define two different searches of the cross-linked condition of the FOX1-FBE data. One search used the restricted set of modifications as published with the first application of CLIR-MS(37), and the other used the expanded set of RNA neutral loss products proposed in this work (validated in **Table 1**). A comparison of the numbers of protein-RNA XLs obtained, both at XLSM and unique XL level, is shown in **Figure 1c**. Not only did the total number of XLSMs increase with the more inclusive search from 313 to 1171 (sum of both replicates), but the quantity of unique amino acid and RNA sequence combinations also increased from 71 to 115 (sum of both replicates). The additional XLs detected fall around the same amino acid positions on the protein as those detected with the more restricted parameters, suggesting the additional identifications contribute to the robustness of the results.

The expanded XL product set therefore builds on the sample preparation and data acquisition optimizations to further increase the sensitivity of the technique and reduce sample amount requirements. Note that this expanded product list was already used to search the data presented in earlier sections describing sample preparation enhancements.

### Validating CLIR-MS data with published structural models

Having optimized sample preparation and data analysis to achieve greater numbers of XL identifications from each CLIR-MS experiment, we then set out to better characterize the structural distance represented by a protein-RNA cross-link, to aid more faithful incorporation of restraints in structural models. We applied the technique to a broader set of protein-RNA complexes containing the proteins FOX1, MBNL1, PTBP1 and SF3A1, for which prior published structural models derived from established structural techniques exist and compared CLIR-MS results for each complex with the respective published structure. These systems represent different modes of RNA recognition, canonical and non-canonical, different protein sizes, and complexes with affinities in the range of single to hundreds of nM (K_d_). For each canonical RNA-binding protein, the protein is bound to a relatively short RNA sequence (7 or fewer nucleotides). The short RNA mimics the behavior of a section of segmentally labelled RNA in a CLIR-MS experiment. The SL4 RNA in complex with SF3A1 is slightly longer, but additionally facilitates evaluation of CLIR-MS in the context of a fixed RNA loop structure.

PTBP1 has many diverse roles in RNA metabolism, including regulation of splicing activity(67) and translation initiation(68). Solution-state NMR structures structures exist for each of its four RRMs in complex with a short polypyrimidine RNA sequence(5) (PDB IDs: 2AD9, 2ADB, 2ADC). The muscleblind-like (MBNL) proteins also act as splicing regulators, controlling tissue specific alternative splicing by targeting CUG and CCUG RNA repeat sequences as shown with the crystal structure of ZnF 3-4 pair bound to RNA(69). Unlike the PTB proteins, RNA binding is mediated by ZnF domains. An NMR-derived structure also exists for the ZnF 1-2 pair of MBNL1 in complex with a short RNA(70) (PDB ID: 5U9B). Interaction of SL4 with SF3A1 was previously observed during formation of pre-spliceosomal complexes(71). Further characterization of this interaction revealed that the ubiquitin-like (UBL) domain found near the C-terminus of SF3A1 mediates the interaction with U1 snRNA SL4(72). As well as being functionally important for splicing, the SF3A1-SL4 interaction is also structurally significant, given that the UBL domain is not considered to be a canonical RNA-binding domain(1). Published structures exist for the U1 snRNP and the SF3A1 protein components in their unbound states. A high-resolution 3D structure of the interaction between SF3A1-UBL and SL4 of the U1 snRNA was also determined separately, using X-ray crystallography and validated using solution-state NMR spectroscopy, functional assays, and protein-RNA XL-MS(48). CLIR-MS data from the FOX1 protein in complex with the FBE, used earlier for method optimization, is also included for comparison to an existing structure (PDB ID: 2ERR).

Taken together, comparisons of CLIR-MS data from these complexes with their respective previously published structures should facilitate inference of a generalizable “reference” protein-RNA cross-linking distance. The selected canonical RNA-binding proteins have published structures determined by solution-state NMR for comparison with CLIR-MS data (where cross-linking takes place in the solution state). Conversely, the structures of the SF3A1 and MBNL ZnF 3-4 bound to RNA were determined with X-ray crystallography, facilitating evaluation of the consistency in measured cross-linking distance between NMR-derived and X-ray crystallography-derived structures.

### CLIR-MS identifies interaction regions in protein-RNA complexes involving known and non-canonical RNA-binding domains

For our experiments, we used full-length PTBP1 protein, an MBNL1 construct spanning positions 1-269, the aforementioned FOX1 RRM construct, and a UBL domain construct spanning amino acids 702-791 for SF3A1. RNAs corresponding to bound nucleic acid species in the published structures for each complex were synthesized, each with 50% natural isotope abundance and 50% heavy isotope RNA. Schematics of each complex are overlaid in **Figure 2a-d**. The optimized CLIR-MS protocol was applied to each complex, and the XLs identified are shown in **Figure 2a-d**.

**Figure 2:**
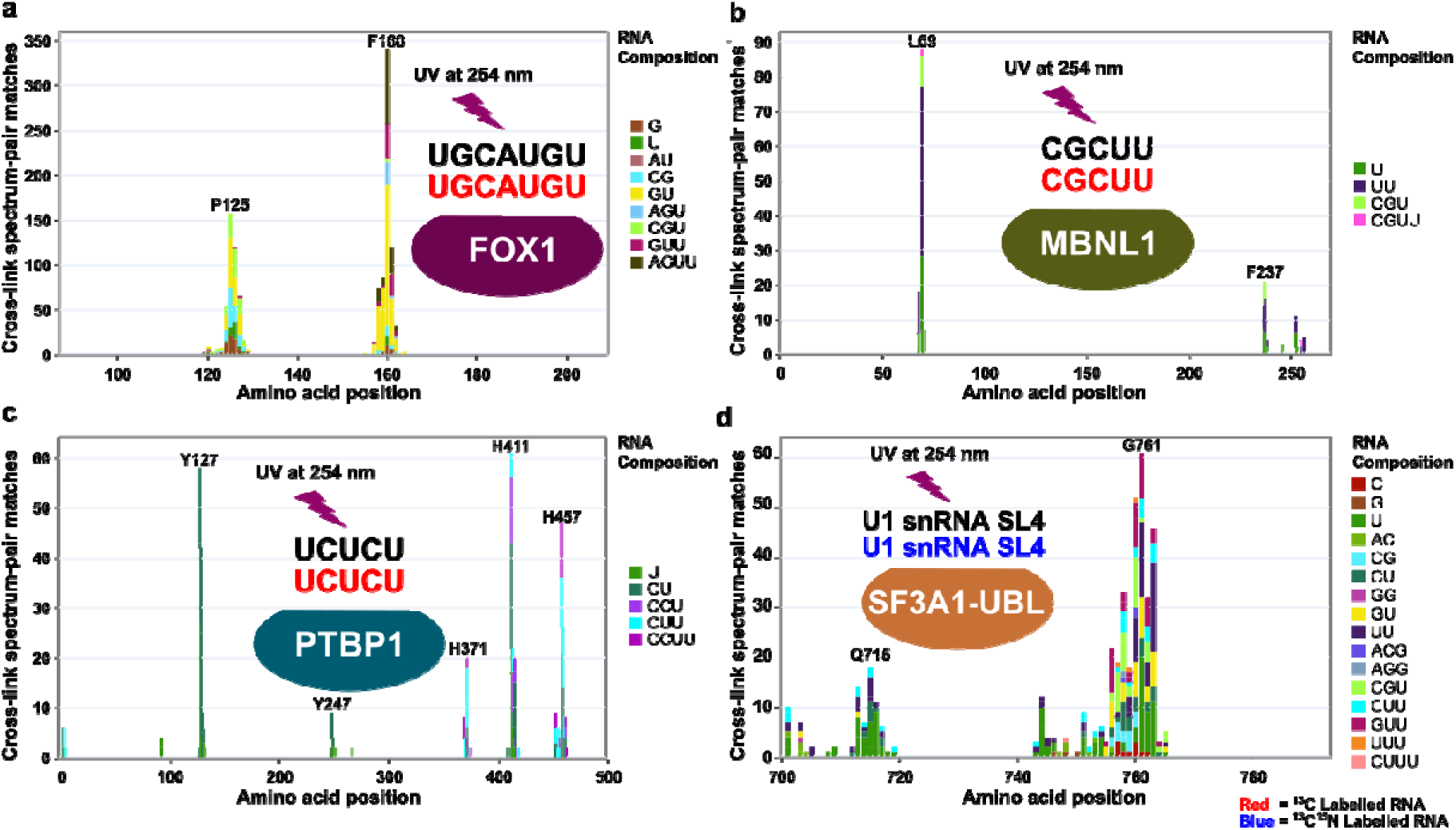
Application of CLIR-MS workflow to model complexes. a)-d) CLIR-MS results (redundant XLSMs) from the complexes FOX1-FBE (stoichiometry protein:RNA = 1:1) a), MBNL1-CGCUU (1:2) b), PTBP1-UCUCU (1:4) c), and SF3A1-UBL with U1 snRNA SL4 (1:1) d), with schematic representation of each protein-RNA complex overlaid.

The optimized sample preparation protocol returned hundreds of XL identifications in each complex,demonstrating the broad applicability of the optimized CLIR-MS protocol to diverse RNA binding proteins. Here, as in previous work, we considered RNA adducts with up to four nucleotides in length; the length distribution varies between samples, probably depending on the nuclease digestion efficiency in the specific samples. While the present section mainly covers the distribution of RNA-binding sites along the primary protein sequences, the following sections will also discuss their agreement with known structures. In the case of the FOX1 RRM, most cross-linked amino acid sites on the protein (**Figure 2a**) fell in clusters around two phenylalanine residues (F126 and F160 respectively). These XLs are well explained by the published structure, with the cross-linked amino acids located on the β-sheet surface of the protein expected to recognize the RNA. Furthermore, these aromatic residues are found π-π stacked with nucleotide bases in the structure, suggesting a close interaction between protein and RNA which may be particularly conducive to cross-linking, as noted elsewhere(40).

In the MBNL1 complex (**Figure 2b**), CLIR-MS analysis suggests that two main groups of amino acids interact with the short RNA (5’-CGCUU-3’), around L69 in ZnF 2 of the 1-2 pair, and around F237 in ZnF 4 of the ZnF 3-4 pair, respectively. This matches published data from MBNL1 interacting with the intronic binding site in human cardiac troponin T pre-mRNA, which demonstrated that only ZnF 2 and ZnF 4 are involved in RNA binding(70). The cross-linking sites in the ZnF 1-2 are well explained by the published NMR-derived structure of RNA interaction with this pair, falling on the position of the protein surface expected to mediate RNA interaction. Prior comparison of unbound ZnF 1-2 and ZnF 3-4 pairs suggests they are highly conserved, both in terms of sequence and structure(70). A crystal structure published earlier also validates the interaction of F237 with RNA(73). The highly cross-linked residues detected by CLIR-MS, L69 (ZnF 2) and F237 (ZnF 4) are also observed at equivalent positions within the tandem ZnFs, suggesting similar structural modes of RNA binding between the two ZnF pairs in this sample.

PTBP1 exhibited fewer amino acid positions cross-linked to RNA when in complex with the short RNA (5’-UCUCU-3‘, **Figure 2c**), than when in complex with a longer RNA such as the IRES RNA (as in previous work by our lab). One potential explanation for this result could be that in the case of the PTBP1-IRES complex, some amino acid sites actively recognize RNA, whereas others make weak unspecific – or as yet uncharacterized – contacts with RNA in the context of a longer RNA bound across multiple RRMs. The latter interactions will be consequently unidentifiable when a shorter RNA is bound, such as the case in **Figure 2**. As with the FOX1 complex, the majority of cross-linking sites are well explained by the published structure (5), falling on the expected exposed β-sheet faces of the proteins across all RRMs, or otherwise nearby on the surface of the protein. The exceptions were in RRM2, where some cross-linking sites were unexpectedly found on the opposite face of the protein than expected. This may originate from a non-specific RNA-protein interaction that may be particularly conducive to UV cross-linking.

Identified XLs from the SF3A1-SL4 complex (**Figure 2d**) highlight two major cross-linking regions in the protein sequence responsible for recognition of the RNA, corresponding to clusters at Q715-K717 (in the β1-β2 loop, UniProt numbering) and at E760-F763 (around strands β3 and β4). The separately described crystal structure(48) revealed an interaction between the C-terminal residues (RGGR motif) with the major groove of the RNA. However, the CLIR-MS analysis, conducted using the same protein construct, yielded no XLs in this region (around amino position 790). This is likely due to inherent incompatibility of the C-terminal sequence, RGGRKK, with trypsin digestion and analysis by LC-MS/MS, with these positions remaining undetectable even in LC-MS/MS analysis of non-cross-linked protein subjected to shotgun proteomics analysis. Distinctively, XLs detected by CLIR-MS (**Figure 2d**) indicate proximity of the β1−β2 loop to the RNA - this interaction is less apparent in the crystal structure, but was in agreement with NMR chemical perturbations and by mutative functional assay, which demonstrate that K717 forms a salt bridge with U1-SL4(48).

For every cross-linked amino acid position detected, unique ribonucleotide compositions at that position were identified, and systematically overlaid to provide a probabilistic representation of interaction sites. Results were plotted as a heat map (data not shown), revealing probable RNA sequence positions with which the respective amino acid sites interact. The analysis revealed that the (G)UUCG(C) terminal loop, inferred from a published model containing SL4(57), cross-links with amino acids around the E760 protein site. However, based on the representation of CLIR-MS XLs in **Figure 2d** alone, some ambiguity remained as to precisely which nucleotides cross-link with the amino acids around Q715.

In summary, our optimized CLIR-MS pipeline identifies (1) multiple RNA-binding regions per protein in the four selected model complexes and (2) the nucleotide composition of RNA regions in contact with proteins through a diverse range of RNA remnants. Moreover, these results are in agreement with benchmark structures from NMR spectrometry, which we used as a reference to capture solution-phase dynamics of these complexes.

### Measuring the distance represented by a protein-RNA cross-link

XL-MS data is frequently used to define distance restraints in structural modelling pipelines, where the distance represented by a XL depends on the reaction chemistry. UV-induced protein-RNA XLs are often referred to as “zero-length” XLs, however in the absence of either a chemical cross-linking reagent or detailed understanding of the chemical structure of the protein-RNA cross-linked species, there is no clear community consensus on the distance over which cross-linking can occur. Empirical comparisons of CLIR-MS data with published structures therefore provide a strategy to understand the distance represented by a UV-induced protein-RNA cross-link, which is vital for faithful use of CLIR-MS data in structural applications.

To gain a better understanding of our data, we filtered XLs identified from each complex for mononucleotide adducts, for which the cross-linked nucleotide position is unequivocal. For each XL we measured the distance from Cα of the amino acid to N1 (for pyrimidines) or N9 (for purines) of the nearest matching nucleotide (backbone to backbone) in the respective published NMR-derived structural model ensembles. (In structural proteomics, backbone-backbone distances are commonly used for modelling, in absence of known side-chain orientations.) The XLs used in each comparison were annotated on the respective published structures, as shown in **Figure 3a-d**. Furthermore, a control set of distances was generated from each of these structures, also from Cα to N1 or N9, but covering all theoretical pairwise combinations of nucleotides and amino acids present in each structural ensemble. The distributions of measured and control distances were plotted for each structure (**Figure 3e-g**). In each case, experimentally detected XL distances form a distribution centered on a shorter distance than, and clearly separated from the theoretical control distances, demonstrating that the XLs capture spatial proximity. The mean protein-RNA cross-linking distances for each sample were 9.7 Å, 10.9 Å, 12.1 Å, and 15.8 Å for the FOX1, PTBP1, MBNL1 and SF3A1 complexes, respectively, with an upper limit of around 20 Å. These restraints are substantially shorter than the size of even the smallest domains of the model proteins used here (compare **Figure 3**).

**Figure 3:**
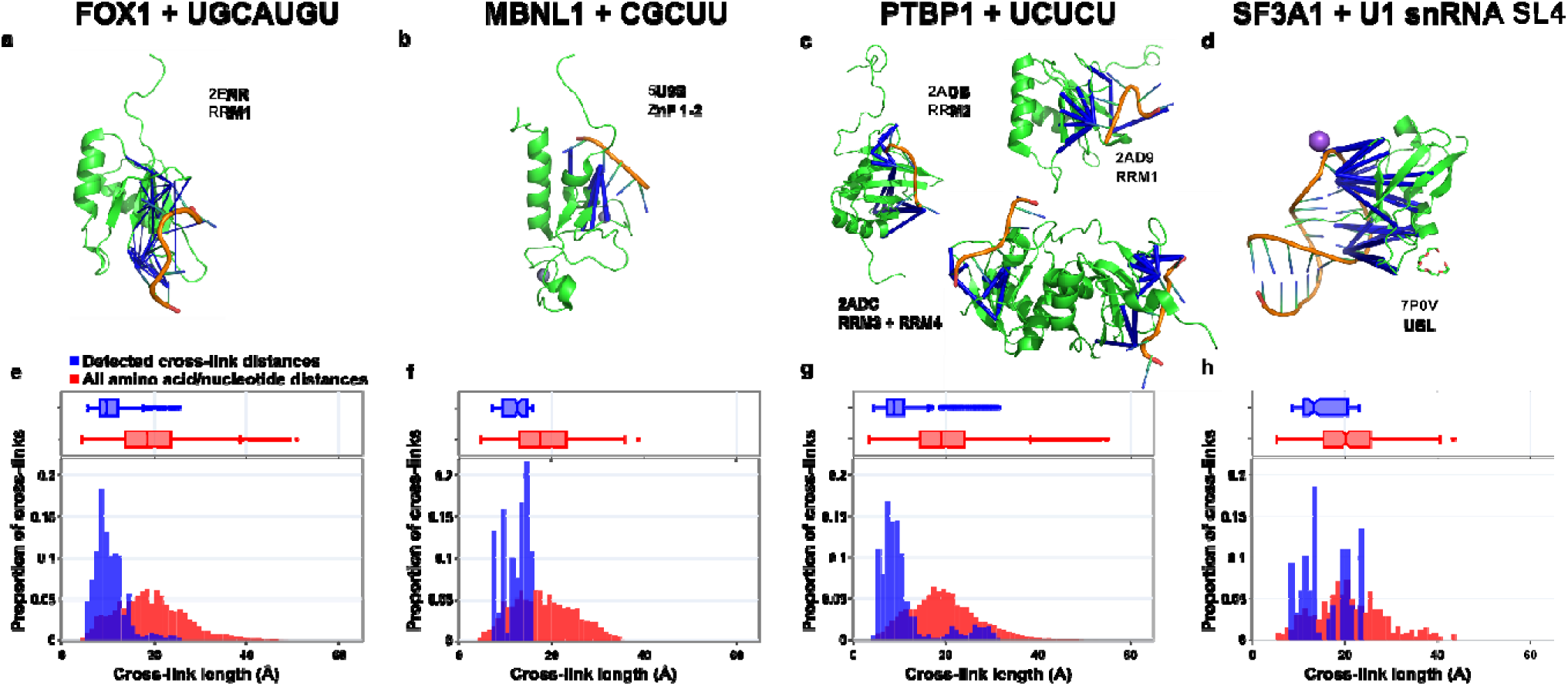
Measuring model complex protein-RNA XL distances against published structures. a)-d) Published structures from the PDB that correspond to the complexes illustrated in Figure 2. XLs involving mononucleotide RNA adducts identified using CLIR-MS are superimposed on the structures in blue. Structures were visualized with PyMOL and the PyXlinkViewer plugin. e)-h) Distribution of distances when XLs from CLIR-MS are measured against the published structures in panels. a)-d), compared with all theoretically possible pairwise combinations of nucleotide and amino acid in each structure. Euclidean distances are measured from the Cα atom of the amino acid to the glycosidic nitrogen atom in the nucleotide (N1 for pyrimidines or N9 for purines). Boxes span quartiles 1-3, with center line representing the median. Whiskers represent upper and lower fences of data points, or highest/lowest values where all values are within this range. e) Measured distances (blue), n=990; Control distances (red), n=18480. f) Measured distances, n=120; Control distances, n=9200. g) Measured distances, n=720; Control distances, n=51960. h) Measured distances, n=123; Control distances, n=359.

A secondary distribution of XL distances greater than 20 Å was observed in both the PTBP1 RRM and in the non-canonical SF3A1 UBL comparison distributions. In the case of the PTBP1 complex, these values derive from XLs in RRM2 which, as mentioned above, may result from comparing CLIR-MS results from a full-length protein with isolated RRM models, rather than true cross-linking distances. In the case of the SF3A1 complex, the longer cross-linking distances result from a relatively flexible linker region, with the longer distances likely representing an artifact of crystallization. By excluding the XLs formed with this region (amino acids 713-719), the mean distance is reduced to 13.6 Å – closer to the mean distances observed with the canonical RNA binding domains.

Whilst observed distances were broadly consistent between the complexes of the canonical RNA-binding proteins, the ZnF-mediated RNA binding of MBNL1 appeared to exhibit slightly longer distances than the RRM mediated binding of PTBP1 and FOX1. This could be explained by differing amino acid compositions of ZnFs and RRMs. The set of all cross-linked amino acid types found in the MBNL1 complex tend to have longer side chains, such as tyrosine, tryptophan, and phenylalanine (although L69 was identified as cross-linked in the most XLSMs). Although these were also found cross-linked in the PTBP1 and FOX1 complexes, XLs with amino acids bearing shorter side chains such as glycine, serine, threonine, and proline were also observed, which may explain the slight shift in distance distributions.

From these comparisons of CLIR-MS derived XLs with prior structural models, we conclude that the XLs detected in a CLIR-MS experiment are depend upon their structural context, and represent a mean proximity of respective peptide and RNA backbones of between 9.7 and 13.6 Å. A certain ambiguity remains as the exact location of the cross-linking sites on the amino acid side chain and the nucleobase remains unknown, and different positional isomers may actually be formed.

### CLIR-MS describes the spatial organization of RNPs in absence of high-resolution information

Protein-protein XL-MS data are frequently employed as a standalone data type for low-resolution placement of proteins relative to one another in a complex(74, 75). In a similar fashion, XLs yielded by the CLIR-MS workflow may be precisely localized on both the protein (by peptide fragmentation and MS/MS) and RNA sequences (by selective isotope labelling and overlay of oligonucleotide adducts found at the same amino acid position of the complex. Furthermore, the structural comparison described above revealed the distances represented by these protein-RNA XLs. Taken together, precisely localized CLIR-MS XLs should therefore contain sufficient structural information to tether an RNA to the correct position on the surface of a protein, providing a low-resolution description of how the two molecules interact. To evaluate this use case, we used DisVis(58, 59) to visualize the accessible interaction space of RNA relative to its corresponding protein in a complex, as constrained by CLIR-MS identified XLs. DisVis can help to classify sets of restraints as mutually supportive or exclusive (e.g., if they represent different interaction sites). It provides as output a density corresponding to the location of an interacting chain (here, the RNA) relative to a stable chain (here, the protein), with the location represented as the center-of-mass of the interactor, compatible with a given number of restraints.

For the canonical protein-RNA complexes, we separated the protein and RNA chains of the published structural models shown in **Figure 3a-c**. We collated a list comprising only mononucleotide XLs that were identified for each complex in the CLIR-MS experiments shown in **Figure 2a-c**. The RNA position associated with each mononucleotide was assumed to be the nearest nucleotide in the published structures, as measured in **Figure 3e-g**. These XLs were specified as restraints with distances from 0 to 12 Å, the upper value in line with the upper quartile of all measured distances (from Cα to N1 for pyrimidines or N9 for purines) observed in **Figure 3e-g**. We then submitted the components to DisVis(59) for occupancy analysis, with protein as the fixed chain and RNA specified as the scanning chain. The outputs are shown in **Figure 4a-c**, where grey shading represents spatial occupancy of the RNA chain relative to the protein (displayed as center-of-mass of the RNA), given the specified XLs. The representation in this figure corresponds to the conformational space compatible with the largest number of restraints that restrict the solution space to ≤0.01% of the conformations sampled, except for MBNL1 (b, 0.39%) and PTBP1 RRM2 (c, 0.07%) where this threshold was too stringent to output any permitted occupancy space. For all canonical protein-RNA complexes, the compatible positioning of the RNA relative to the protein derived from the CLIR-MS XLs closely resembled the placement of the RNA in published structural models of each of these complexes (**Figure 3a-c**), and only one preferred interaction region was observed in all cases. As one might expect, the permitted occupancy space is smaller when a greater number of XLs are used to define the space (data not shown). Conversely, requiring fewer restraints to be fulfilled at the same time may result in additional contact regions emerging, as is seen for RRM2 in the PTBP1-UCUCU complex (data not shown).

**Figure 4:**
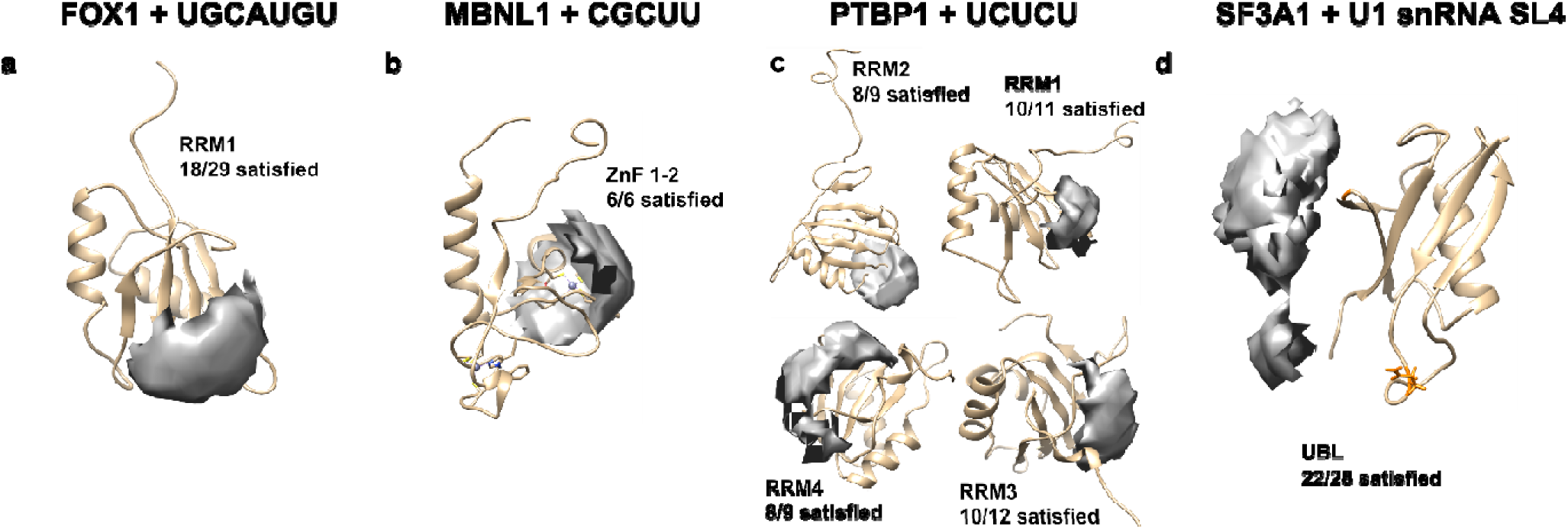
Visualizing the structural information contained in CLIR-MS protein-RNA XLs when used as an independent data type. a)-d) Visualization of the structural information contained in point-to-point distance restraints obtained from data in Figure 2a-d using DisVis. Visualizations are produced using UCSF Chimera.

In each of the relatively small canonical protein-RNA complexes, RNA contact sites fall close together on the surface of the protein. Employing short RNA sequences, it is sometimes challenging to precisely determine RNA orientation due to a large degree of rotational freedom for the RNA. However, cross-links detected from the longer structured RNA of the non-canonical SF3A1-UBL-SL4 complex could provide enough information for more precise orientation based on CLIR-MS cross-links alone. To answer with high confidence which nucleotides in SL4 interact with amino acid Q715 (**Figure 2d**), DisVis analysis was performed, using previously published standalone protein and RNA structures (PDB ID: 1ZKH for the unbound SF3A1 UBL structure(56), and PDB ID: 6QX9 for the structured SL4 RNA(57), to predict a set of mutually spatially compatible XLs. We used the highest scoring 5% of protein-RNA contact site position pairs as distance restraints.

As with the canonical protein-RNA complexes, a permitted occupation space of the SL4 RNA relative to the SF3A1-UBL protein was calculated with DisVis, based upon a subset of the ambiguous distance restraints (**Figure 4d**). DisVis assigns a *z*-score to each specified restraint to determine which of the tested restraints were most frequently violated during the occupancy analysis. The XLs with the most favorable (lowest) *z*-scores are located between the nucleotides at the top of the RNA stem loop structure, and amino acids E760-F763 in the protein, consistent with the probabilistic analysis. Based on these analyses, the UUCG tetraloop contacts the surface of the protein near E760-F763. The other major cross-linked amino acid site suggests that the lower part of the stem loop is then tethered to the protein around Q715-K717. Unlike with the canonical protein-RNA complexes in **Figure 4a-c**, the contact sites between protein and RNA are more spatially separated on the protein surface, suggesting that there is directionality to the permitted occupancy space of the RNA. Comparison of the directional occupation space shown in **Figure 4d** with the *bona fide* high resolution structure of this interaction (described separately(48)) validated the predicted position of the RNA relative to the protein, derived here using only CLIR-MS restraints.

Taken together, our findings confirm that CLIR-MS distance restraints are generally sufficient as a standalone low-resolution descriptor of spatial RNA occupancy relative to an RNA binding protein. Furthermore, in cases with long structured RNAs, CLIR-MS restraints may facilitate description of both the position and the orientation RNA structure in relation to RNA binding proteins. Characterizations carried out using this methodology therefore represent a reliable starting point for sophisticated, atomic-scale integrative structural modelling workflows(76).

### Expansion of the CLIR-MS concept to 4-thio-uridine labeled RNA

Due to the inherently low reaction yields of protein-RNA cross-linking reactions, many experimental workflows that rely on UV cross-linking of protein to RNA substitute uracil for 4-thio-uracil (4SU) to increase the proportion of protein-RNA complex that is cross-linked(38, 64, 77, 78). For example, 4SU-labeled enhancer RNAs were shown to provide higher XL site coverage on an RNA polymerase II-DSIF-NELF complex(78). We postulated that 4SU could also be used with CLIR-MS to increase the reaction yield, and hence the ability to detect XLs. However, little is known about the cross-linking preferences and distance restraints of 4SU-assisted crosslinking, limiting its potential in structural application.

Production of RNA by solid phase synthesis enables position-specific incorporation of chemically modified nucleotides such as 4SU, meaning the cross-linking behavior of 4SU at a specified nucleotide position can be evaluated. Three separate FOX1-FBE samples were prepared, each with one of the three uracil positions in the FBE RNA heptanucleotide replaced with 4SU (schematics overlaid in **Figure 5a**). Samples were irradiated with 365 nm UV light to ensure that the XLs formed only at the 4SU-substituted base. Samples were then analyzed using the optimized CLIR-MS workflow and identified XLs are shown in (**Figure 5a**). As expected, the numbers of XLSMs are relatively high, compared with a similar sample mass using natural nucleotides shown in previous figures, and especially so, considering that all XLs derive from a single nucleotide position. Most cross-linking reactions occurred at positions U1 and U7. According to the published structure, U1 exhibits some conformational heterogeneity in its binding, and U7 is not held rigidly in place by specific hydrogen bonds at its Watson-Crick face(43). Position U5, which the published structure indicates is firmly held in place by hydrogen bonds to multiple amino acid residues(43), did not XL so strongly. The major cross-linked protein sites differed from those obtained in samples containing only natural nucleotides, with a loss of XLs around amino acid position 160 when any of the uracil positions were replaced with 4SU, and a gain of XLs surrounding N151. However, these amino acid positions are still close to RNA in the published structure.

**Figure 5:**
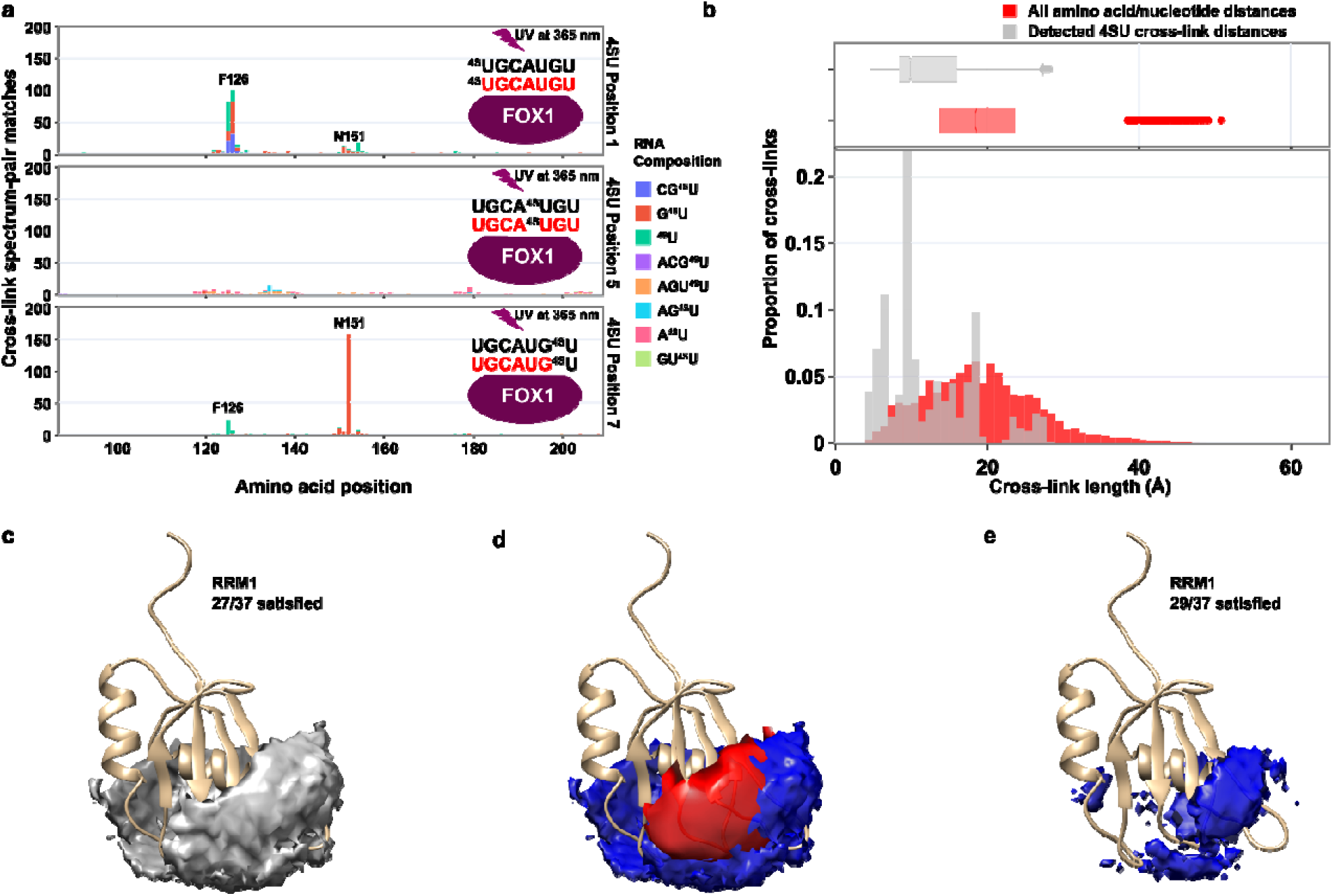
4-thio-uracil leads to qualitative as well as quantitative changes to results compared with natural uracil. a) CLIR-MS results for FOX1-FBE after replacing specific uracil positions with photoactive 4-thio-uracil. 6 XLSMs across all label position conditions that were identified as not containing 4SU (likely mis-assignment due to isobaric mass) are excluded. b) XL distances detected in the FOX1-FBE complex where 4-thio-uracil is used in place of natural uracil, as measured against the published structure for the protein-RNA complex. Boxes span Q1-Q3, with center line representing the median. Whiskers represent upper and lower fences of data points, or highest/lowest values where all values are within this range. g) Measured distances (grey), n=4650; Control distances (red), n=18480. (c) DisVis RNA occupancy space permitted by 4SU-derived cross-links with respect to the FOX1 RRM. Occupancy space shown according to the same cut-off used in Figure 4, with the solution space restricted to ≤0.01% of the conformations sampled. (d) Overlay of the occupancy space from 4SU-derived cross-links at ≤0.01% of sampled conformations (blue) with the equivalent space from canoncial nucleobase-derived cross-links (red, from Fig. 4a). (e) 4SU-derived occupancy space with an increased number of satisfied restraints such that the volume of the permitted space is approximately equal to that from canoncial nucleobase derived cross-links.

DisVis occupancy analysis was once again carried out with the FOX1 protein and FBE RNA, this time using the 4SU-derived cross-links as distance restraints. The resulting space is visualised in Figure 5c; the data lead to a larger theoretical solution space, apparently with a greater number of conformations sampled. Therefore, when plotted according to the same thresholds in Figure 4 (lowest number of XLs required to restrict the solution space to ≤0.01% of the conformations sampled), the RNA is less precisely localised than with XLs derived from canonical nucleobases. The canonical and 4SU occupancy spaces are overlaid in Figure 5d, with volumes of 687 and 2719 arbitrary units, respectively. By increasing the number of satisfied restraints to 29, the volume of the 4SU-based occupancy space can be approximately reduced to the same as that of the canonical nucleobase analysis. The reduced space is shown in Figure 5e has a volume of 655 arbitrary units. Compared with the canonical nucleobase analysis shown in Figure 4a, the RNA is less precisely located with respect to the beta-strand face expected from the published NMR-derived structure. Rather, the permitted occupancy space stretches around to the alpha helix on the opposite face of the protein. Taken together, these results indicate that whilst 4SU-derived cross-links may provide structurally valid information, the nature of reactivity may differ substantially from canonical nucleobases.

Next, we compared identified 4SU-derived XLs with the published FOX1-FBE structure ensemble, with distances measured from Cα of the amino acid to N1 of 4SU. The distribution of observed distances and the control distance set containing all possible amino acid and nucleotide pairs in the complex are shown in **Figure 5b**. The distribution of 4SU XLs is well resolved from the control set of distances, similar to the canonical uracil XLs shown in **Figure 3d**, with a mean distance of 12.2 Å for the measured distances. Taken together, despite the positionally distinct cross-linking observed in **Figure 5a** with respect to canonical uracil, the cross-linking distance represented by a 4SU-derived XL appears similar to canonical uracil.

## Discussion

The enhancements to the CLIR-MS protocol presented here provide a much higher greater coverage of the UV cross-linked species created in a sample compared to that obtained following the originally described protocol. This major improvement derives from a distribution of RNA modifications over a set of consecutive amino acids, rather than linking at a single amino acid position. The increased density of XLs now identified in each sample raises confidence in a detected protein-RNA interaction site, increasing the standalone value of CLIR-MS data.

The variety of RNA-derived peptide modifications observed is striking. The analysis approach used here considers the “biological” information contained in a XL (i.e. protein proximity to a given nucleotide sequence) constant between neutral loss products (**Table 1**) with the same RNA sequence. Whilst the biological information contained in different neutral loss products with the same sequence composition is equal, this diversity may have implications for the analytical workflow. Despite the enrichment step, peptide-RNA adducts are often present in very low in abundance in the final LC-MS/MS sample, close to the limits of detection. If the variety of RNA-derived adducts depends upon sample preparation, further optimizations could be considered to reduce the number of adduct types produced. For example, different combinations of nucleases may leave distinct RNA adduct types; of the nucleases used here, RNases A(79) and T1(80) yield a 5’ hydroxy product, but leave a 2’-3’ cyclic phosphate or a 2’ or 3’ phosphate attached (which may explain the observed -H_2_O loss). However, benzonase leaves the phosphate attached(81), as noted previously(38). Such technical stratification may unnecessarily reduce signal intensity. Alternative approaches using chemical cleavage of RNA in comparable experimental setups have been recently demonstrated(40, 82), which may reduce the variety of RNA product types, but at the same time result in near-exclusively mononucleotide attachments to peptides. The reduced proportion of polynucleotide adducts resulting from this approach may however make it more challenging to precisely assign the cross-linking site on an RNA sequence, because the data lack the required sequence context on the RNA side.

To be useful as distance restraints, protein-RNA XLs should be of a well-defined length and localized both to a single amino acid on the peptide, and to a single nucleotide on the RNA. In experiments shown here, localization on a peptide was achieved by MS/MS, analagous to PTMs in conventional MS-mediated proteomics experiments. For the RNA side, short stretches of segmentally labelled RNA used in a CLIR-MS experiment narrow the cross-linked ribonucleotides to those within the labelled sequence, as isotope pairing is a requirement to produce an identification. Some further analysis is however required to refine the position to a single nucleotide. As shown previously(37), this may be achieved by overlaying detected mono-, di-, and trinucleotide species. This approach was applied systematically such that probable sites of RNA interaction are computed for every cross-linked amino acid position. This “heatmap” feature is available in the recently introduced RNxQuest package(54). The rich variety of polynucleotide RNA sequence compositions found linked to a particular peptide therefore together contain the information to localize the XL at up to single nucleotide resolution. We therefore consider them beneficial enough in structural studies to select polynucleotide adduct-yielding nuclease digestion over alternative chemical RNA degradation approaches(82).

Comparing published structures with protein-RNA cross-linking data from CLIR-MS experiments performed on different types of RNA binding domain demonstrated that UV-induced protein-RNA XLs consistently form over a mean distance of 9.7-13.6 Å (measured backbone to backbone). A clear understanding of this parameter is vital for structural interpretation, if cross-linking data is to be used to specify restraints that reflect true proximity. The distance measurements shown here appear to agree with the chemical structures of cross-linking products proposed in prior literature, which suggests a mechanism for UV-induced protein-RNA cross-linking(39, 83).

Incorporation of 4SU in place of uracil is conventionally accepted by the scientific community as a strategy for increasing the yield of a protein-RNA UV cross-linking experiment, under the assumption that structure or function of an RNA are minimally impacted(38, 77, 78, 84). The data presented here may subtly alter how such data is interpreted in a structural context. The observation that canonical nucleobases and 4SU have similar measured cross-linking distances in known structures supports the assumption that this replacement does not have structural consequences. However, the positional distinction shown at the level of amino acid cross-linking sites between the two nucleobase types may warrant more caution when choosing this approach. Recently published proteome-wide studies of RNA binding proteins captured by RNA pull-down also highlight distinct behaviors of uracil and 4SU, with each pulling down a different subset of the proteome(63). Our observation that 4SU induced XLs lead to distinct cross-linked amino acid positions of FOX1 compared with natural uracil is consistent with 4SU cross-linking capturing a distinct subset of protein-RNA binding interactions from natural bases in the proteome-wide study. We speculate that the differences may result from distinct lifetimes of the radical species generated when a natural nucleotide is irradiated compared with a 4SU. Indeed, more fundamental studies of sulphur-substituted nucleotides provide evidence of a longer-lived triplet state radical(85), although further experimental work would be required to examine this hypothesis. Taken together with the results presented here, these observations suggest that 4SU and natural uracil may have subtly distinct behaviors beyond the difference in reaction yields, which should be carefully considered when interpreting results generated using 4SU cross-linking.

The DisVis analyses of protein-RNA XLs demonstrate clearly the information content of CLIR-MS results. Whilst the RNA binding behavior of many canonical RNA binding protein domains is well understood, a large array of novel RNA-binding protein domains are discovered(1). The structural characterization of every novel RNA binding protein using established structural biology techniques will be an enormous undertaking for the scientific community, hence technical advances that enable the process are an attractive prospect. In the model complexes with prior structures studied here (**Figure 4)**, CLIR-MS derived restraints contain sufficient information to position the RNA on the correct RNA binding surface of the protein, even in the absence of other complementary data types. This represents an additional use case to the one shown previously(37), where an integrative modelling approach used multiple structural data sources to determine a final model.

The protein-RNA complexes studied in **Figure 4a-c** all have short, single stranded and flexible RNAs, and in all, especially in the case of the MBNL1 complex, cross-linked amino acids tend to fall within a small spatial cluster on the surface of each protein. RNA occupation spaces provided by DisVis therefore reflect the remaining rotational degrees of freedom. In the case of the SF3A1-UBL interaction with U1 snRNA SL4 shown in **Figure 4d**, the RNA instead forms a rigid stem-loop structure, and therefore, multiple clusters of cross-linked amino acids are spread more widely across the surface of the protein. This results in a narrower and more elongated occupation space. These differing behaviors could indicate that CLIR-MS data is most successful as a standalone data type when applied to study interactions of conformationally rigid RNA structural features with a protein, and where multiple, spatially separated protein-RNA contact sites are identified by cross-linking.

Overall, the optimized CLIR-MS protocol and data analysis approach provide much greater numbers of identifications than the original protocol(37), improving the confidence in identified protein-RNA interaction sites detected. These identifications compare favorably with existing structures derived from other established structural techniques, when used to study complexes with well characterized structures. We used these comparisons to make general inferences about the distances over which UV-induced protein-RNA XLs form, and the robustness of protein-RNA cross-linking data. Our optimizations and observations guide the interpretation of protein-RNA cross-linking data, whilst demonstrating a new use case as an independent source of structural data. We therefore propose that CLIR-MS data is well suited to low-resolution binding interface characterization for rigid complexes when considered in isolation, or as an additional complementary data type in more sophisticated integrative modelling pipelines(76) to achieve high-resolution, atomic-scale models. In cases of the latter, CLIR-MS data may prove particularly valuable when probing flexible/unstructured protein regions where more established structural techniques relying on conformational homogeneity may struggle to provide coverage, as shown in our previous work on PTBP1 (37,47) and the SARS-CoV-2 nucleocapsid protein (66), where we combined cross-linking data with NMR spectroscopy, electron paramagnetic resonance and small-angle scattering. CLIR-MS does not distinguish between binding to structured and unstructured regions as long as no amino acid sequences that are difficult to probe by mass spectrometry are present (e.g., for enzymatic digestion). In such cases, proteases other than trypsin can be employed to produce peptides of appropriate length.Such integrative approaches remain highly relevant as even the most recent versions of deep learning-based structure prediction tools such as AlphaFold 3 perform poorly on protein-RNA complexes. In summary, our method may prove a useful tool to reliably study the emerging plethora of non-canonical RNA binding domains with the relative speed of an MS-based pipeline.

## Data Availability

Proteomics raw data and xQuest outputs have been deposited to the ProteomeXchange Consortium (http://proteomecentral.proteomexchange.org) via the PRIDE partner repository with the dataset identifier PXD029930. CLIR-MS results shown for the SF3A1 complex were generated by reanalysis of a dataset described elsewhere(48), and are deposited in the PRIDE repository with dataset identifier PXD027189.

Scripts used to measure distances in structural models have been deposited in the ETH Research Collection and are available via the DOI 10.3929/ethz-b-000683144.

## Acknowledgements

We thank P. Picotti for access to laboratory and MS infrastructure. We thank C. von Schroetter for assistance in producing *in vitro* transcribed RNA. We thank the lab of C. Branlant for the MBNL1 plasmid. We thank N. de Souza for critical comments during preparation of the manuscript.

## Author Contributions

A.L., F.H.-T.A., J.H. and R.A. conceived the study, and supervised C.P.S., A.K. and T.d.V. A.K. produced chemically synthesized RNA. T.d.V. produced double-isotope labelled RNA. A.K. and T.d.V. (FOX1), T.d.V. (SF3A1, PTBP1, MBNL1), I.B. (MBNL1) expressed and purified proteins for the study. A.K. and T.d.V. performed cross-linking of protein-RNA complexes. C.P.S. performed TiO_2_ enrichment, C_18_ clean-up, and measured samples by mass spectrometry. C.P.S. and M.G. wrote analysis scripts and conducted data analysis using xQuest. C.P.S. performed DisVis analysis. C.P.S. and A.L. wrote the manuscript together. All authors contributed to manuscript revisions and approved the final manuscript.

## Notes

Present address: Chris P. Sarnowski, DISCO Pharmaceuticals Swiss GmbH, Wagistrasse 25, Schlieren, Switzerland

Present address: Anna Knörlein, Chemical Biology Program, Memorial Sloan Kettering Cancer Center, New York City, NY, USA

Present address: Michael Götze, Department of Biology, Chemistry, Pharmacy, Institute of Chemistry and Biochemistry, Freie Universität Berlin, Berlin, Germany

Present address: Irene Beusch, Institute of Molecular Infection Biology, Julius-Maximilians-Universität Würzburg, Würzburg, Germany

